# Disentangling Context-Specific Alzheimer’s Disease Progression Signatures through an Explainable AI Framework

**DOI:** 10.64898/2026.11.02.763789

**Authors:** Allison W. Li, Wei Qiu, Chris Arian, Jessica E. Young, Su-In Lee

## Abstract

Alzheimer’s disease (AD) progression involves stage-dependent transcriptional changes that are often masked by dominant cellular and biological variation. Here, we present **ADVISE (Alzheimer’s Disease Variational Isolation of Signal Embeddings)**, an interpretable deep generative framework designed to disentangle Braak stage-associated transcriptional signals from background biological variation through separate latent representations. Using explainable AI, ADVISE prioritizes genes and pathways contributing to the disease-specific representation. Applying ADVISE to large-scale single-nucleus RNA sequencing data from the human prefrontal cortex, we identify shared and context-specific signatures of AD progression across global, sex-specific, and cell-type-specific analyses, revealing convergent molecular programs involving synaptic transmission, HSF1-associated proteostatic stress responses, and SLIT/ROBO axon guidance. Notably, ADVISE prioritizes *ROBO2* as a progression-associated candidate in excitatory neurons. Experimental validation using CRISPR activation in hiPSC-derived excitatory neurons demonstrates that *ROBO2* activation reduces the phosphorylated Tau-to-total Tau ratio. ADVISE thus reveals context-dependent molecular signatures of AD progression and bridges human single-cell discoveries with experimental validation of candidate regulators of AD pathology.

## Introduction

Alzheimer’s disease (AD) is the leading cause of dementia, accounting for approximately 60–70% of cases worldwide. The global number of people living with dementia is projected to nearly triple, from 57 million in 2019 to 153 million by 2050 [1, 2]. It is a major cause of death and disability, imposing substantial social and economic burdens [2, 3]. Although advances in diagnostic biomarkers have improved our ability to detect and characterize AD pathology, the molecular mechanisms underlying progression from early to advanced disease stages remain incompletely understood.

A central obstacle is that AD progression is neither linear nor uniform. Rather than following a simple pathological cascade, AD involves dynamic interactions among amyloid accumulation, Tau pathology, neuroinflammation, synaptic dysfunction, and vascular changes [4, 5]. These interacting processes contribute to stage-dependent transcriptional changes that vary across biological contexts. Neurons, microglia, astrocytes, and oligodendrocytes exhibit distinct vulnerabilities and transcriptional responses during progression [6, 7], while sex influences AD risk, pathology, and molecular responses [8, 9, 10, 11]. Understanding AD progression therefore requires distinguishing molecular signatures shared across biological contexts from those specific to individual cell types and sexes.

Single-nucleus RNA sequencing (snRNA-seq) of postmortem human brain tissue provides the cellular resolution needed to address these questions, but extracting progression-associated signals from these data poses substantial challenges. Disease-associated transcriptional variation can be masked by dominant sources of biological and technical variation, including cell type, sex, age, and batch effects. Moreover, conventional case–control comparisons may overlook transcriptional changes associated with intermediate pathological stages, limiting our understanding of how molecular programs vary across the AD spectrum. These challenges are further compounded by heterogeneous disease-associated responses across cell types and between sexes, making it difficult to distinguish shared molecular programs from context-specific signatures. Finally, connecting these transcriptional signatures to biological mechanisms requires identifying the genes and pathways underlying progression-associated variation and prioritizing candidates for experimental validation.

Despite these challenges, numerous studies have applied single-cell and single-nucleus transcriptomics to AD, providing important insights into its cellular and molecular pathology [7, 12, 13, 14, 6, 15, 16, 17]. These studies have identified selectively vulnerable excitatory neuron populations in early-affected regions [6], disease-associated glial states including TREM2-dependent microglial responses [14], cell-type-specific transcriptional changes associated with pathology and cognitive resilience [7, 15], and sex-dependent expression patterns across major cell types [18, 19]. Large multimodal atlases have further begun to characterize variation across the continuum of AD pathology [16, 17]. Although these studies have substantially advanced our understanding of AD-associated molecular changes, systematically distinguishing shared and context-specific transcriptional programs associated with disease progression–and linking these signatures to experimentally testable molecular candidates–remains challenging.

Computational approaches have played a central role in these discoveries, ranging from graph-based unsupervised modeling to statistical modeling and deep representation learning. Graph-based approaches, including nearest-neighbor clustering and graph-based trajectory inference, have enabled the identification of diverse cell populations and disease-associated states [7, 13, 12, 14, 18]. However, these approaches can be influenced by dominant sources of variation unrelated to disease progression, while discrete clustering may obscure gradual transcriptional changes across pathological stages. Statistical approaches, including cell-level and mixed-effects models for differential expression analysis, allow explicit adjustment for biological and technical covariates [7, 14, 15, 18, 17], but commonly used linear formulations may not fully capture nonlinear relationships or interactions among disease stage, cell type, and sex. Deep generative models can capture nonlinear transcriptional variation and account for technical effects [20, 21], and have been applied to AD datasets for data integration and reference mapping [6, 16]. Nevertheless, disentangling progression-associated transcriptional signals from dominant background variation, while preserving context-dependent disease signatures and enabling gene-level interpretation, remains challenging.

To investigate these unresolved biological questions, we developed ADVISE (Alzheimer’s Disease Variational Isolation of Signal Embeddings), an interpretable deep generative framework for identifying transcriptional signatures associated with AD progression. Building on ACE (Aging Cell Embeddings), a previously developed framework for disentangling aging-associated signals from background variation [22], and related approaches to disentangled representation learning [23], ADVISE learns separate latent representations for Braak stage-associated transcriptional variation and background biological and technical variation. By incorporating Braak stage as an ordinal measure of neuropathological severity, ADVISE captures transcriptional variation across pathological stages rather than relying solely on binary diagnostic comparisons. To resolve context-dependent signatures, we employ complementary modeling strategies for global, cell-type-specific, and sex-specific analyses. Finally, explainable AI enables attribution of the Braak stage-associated representation to individual genes, facilitating pathway analysis and prioritization of candidates for experimental investigation.

In this study, we apply ADVISE to large-scale snRNA-seq data from the human prefrontal cortex to characterize molecular signatures associated with AD progression across global, cell-type-specific, and sex-specific contexts. We identify both shared and context-dependent transcriptional programs, implicating synaptic transmission, HSF1-associated proteostatic stress responses, and *SLIT/ROBO* axon guidance. Notably, the *SLIT/ROBO* pathway emerges across multiple analytical contexts, and *ROBO2* is prioritized as a progression-associated candidate in excitatory neurons. Consistent with this association, *ROBO2* expression decreases with increasing Braak stage. To investigate its functional relevance, we performed CRISPR activation of *ROBO2* in human induced pluripotent stem cell-derived excitatory neurons and found that its activation reduces the phosphorylated Tau-to-total Tau ratio. These findings connect context-resolved transcriptional analysis of human AD tissue with experimental evidence implicating *ROBO2* in AD-relevant Tau pathology.

## Results

### ADVISE disentangles AD progression-associated transcriptional signals from background variation

To investigate molecular signatures associated with Alzheimer’s disease (AD) progression across diverse biological contexts, we developed ADVISE, an explainable deep generative framework designed to disentangle Braak stage-associated transcriptional variation from background biological and technical variation in single-cell RNA sequencing data (Figure 1a). ADVISE builds on ACE (Aging Cell Embeddings) [22], a previously developed variational autoencoder-based framework for separating aging-associated transcriptional variation from background signals. To resolve context-dependent AD signatures, we employed distinct modeling strategies for cell-type-specific and sex-specific analyses. We applied ADVISE to the single-nucleus RNA sequencing dataset from Mathys et al. (2019) [7], comprising approximately 80,000 nuclei from the human prefrontal cortex of 48 individuals spanning the AD spectrum.

**Figure 1.**
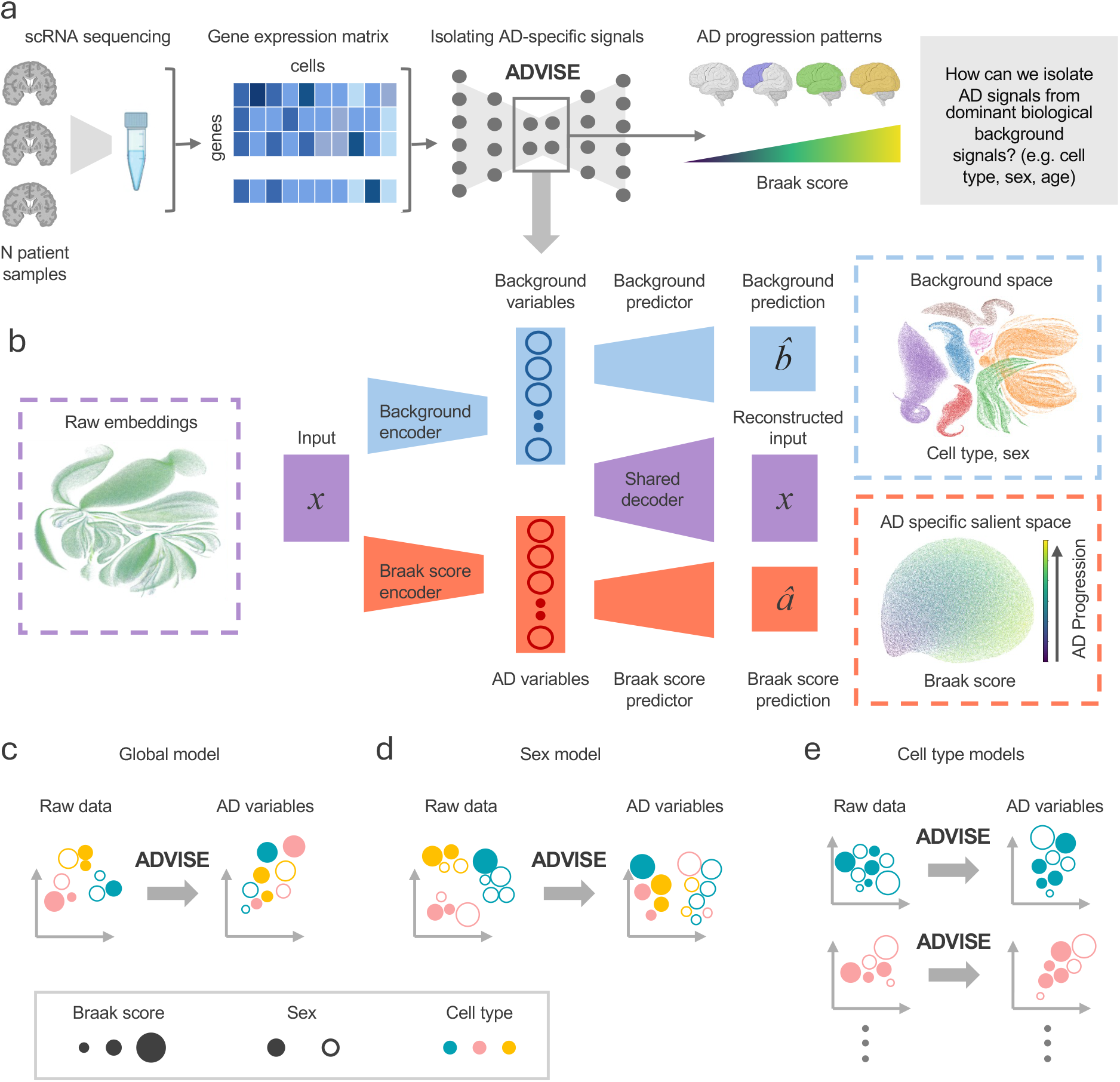
Overview of the ADVISE framework for disentangling AD signatures in single-cell RNA-seq data. **(a)** ADVISE is a framework designed to isolate AD-specific transcriptional signatures from scRNA-seq data by separating AD-related variation from dominant background biological covariates (e.g., cell type, sex). **(b)** ADVISE has two encoders: an AD encoder that captures target AD signals, and a background encoder that captures dominant covariate signals. The extracted AD variables are then used to predict Braak scores to capture AD progression, while the extracted background variables are used to predict covariate labels (e.g., cell type, sex). A shared decoder reconstructs the input from both AD variables and background variables, encouraging each set of variables to capture distinct yet complementary information. **(c, d, e)** ADVISE is implemented with three settings to disentangle global (i.e., shared across covariates) and covariate-related AD signatures (e.g. sex, cell type). Circle size indicates Braak stages intensity; fill denotes sex; color represents cell type. **(c)** A global model is trained to capture global AD signatures across the dataset. **(d)** A sex-specific model is trained to capture sex-related AD signatures. The resulting AD variables reflect gene expression signatures driven by sex-related contributions to AD. **(e)** Cell type-specific models are trained to capture AD signatures specific to each cell type. The resulting AD variables capture gene expression signatures related to AD progression within each cell type.

The ADVISE architecture (Figure 1b; Methods) consists of two parallel encoder networks that learn complementary latent representations. The first encoder learns a salient representation capturing Braak stage-associated transcriptional variation, while the second learns a background representation capturing other sources of biological and technical variation. Auxiliary prediction networks guide this separation: the salient representation is trained to predict Braak stage, an ordinal measure of neurofibrillary tangle pathology, whereas the background representation is trained to predict specified background covariates. To further encourage disentanglement, ADVISE incorporates the Hilbert–Schmidt Independence Criterion (HSIC) [24, 25, 26], which penalizes statistical dependence between the two latent representations. A probabilistic decoder reconstructs observed gene expression counts using the learned representations and the specified conditioning variables. Building on the count-based generative modeling framework of scVI [20], ADVISE accounts for sequencing depth and other technical sources of variation in single-nucleus RNA sequencing data.

AD progression is associated with both molecular signatures shared across biological contexts and context-specific transcriptional changes that differ by sex and cell type [27, 18, 28, 29]. To characterize these shared and context-dependent signatures, we employed three complementary ADVISE modeling configurations: (i) a global model identifying Braak stage-associated transcriptional patterns across all cells, (ii) a sex-specific model characterizing sex-dependent progression-associated signatures, and (iii) a cell-type-specific model resolving progression-associated signatures within individual cell types (Figure 1c-e). For each configuration, we applied explainable AI (XAI) methods to identify and rank genes contributing to the Braak stage-associated representation, followed by pathway analysis to characterize the underlying biological processes.

### ADVISE captures global AD signals shared across sex and cell types

To characterize global AD progression-associated signals, we designed an ADVISE model that explicitly captures Braak score-associated variation that is consistent across sex and cell type (Figure 2a). In this model, ADVISE was trained on the unstratified dataset, with Braak score encoded in the AD latent space and cell type and sex, representing dominant background biological covariates, assigned to the background space. This design enables the model to isolate AD progression signals that are consistent across biological contexts, such that the learned AD variables are largely independent of sex- and cell type-driven variation. We first evaluated the model’s ability to capture Braak score-related variation in the AD space. In the raw dataset, AD-associated progression trends are not readily apparent, with the dominant structure in the data driven primarily by cell type-specific differences (see Supplementary Figure 1 for UMAP visualizations of the unprocessed data). After training, however, ADVISE learned an AD space in which Braak score embeddings form a continuous disease progression trajectory that is largely independent of sex and cell type effects (Figure 2b). This suggests that the model successfully disentangles AD progression from background biological covariates that otherwise obscure disease-related signatures in the data.

**Figure 2.**
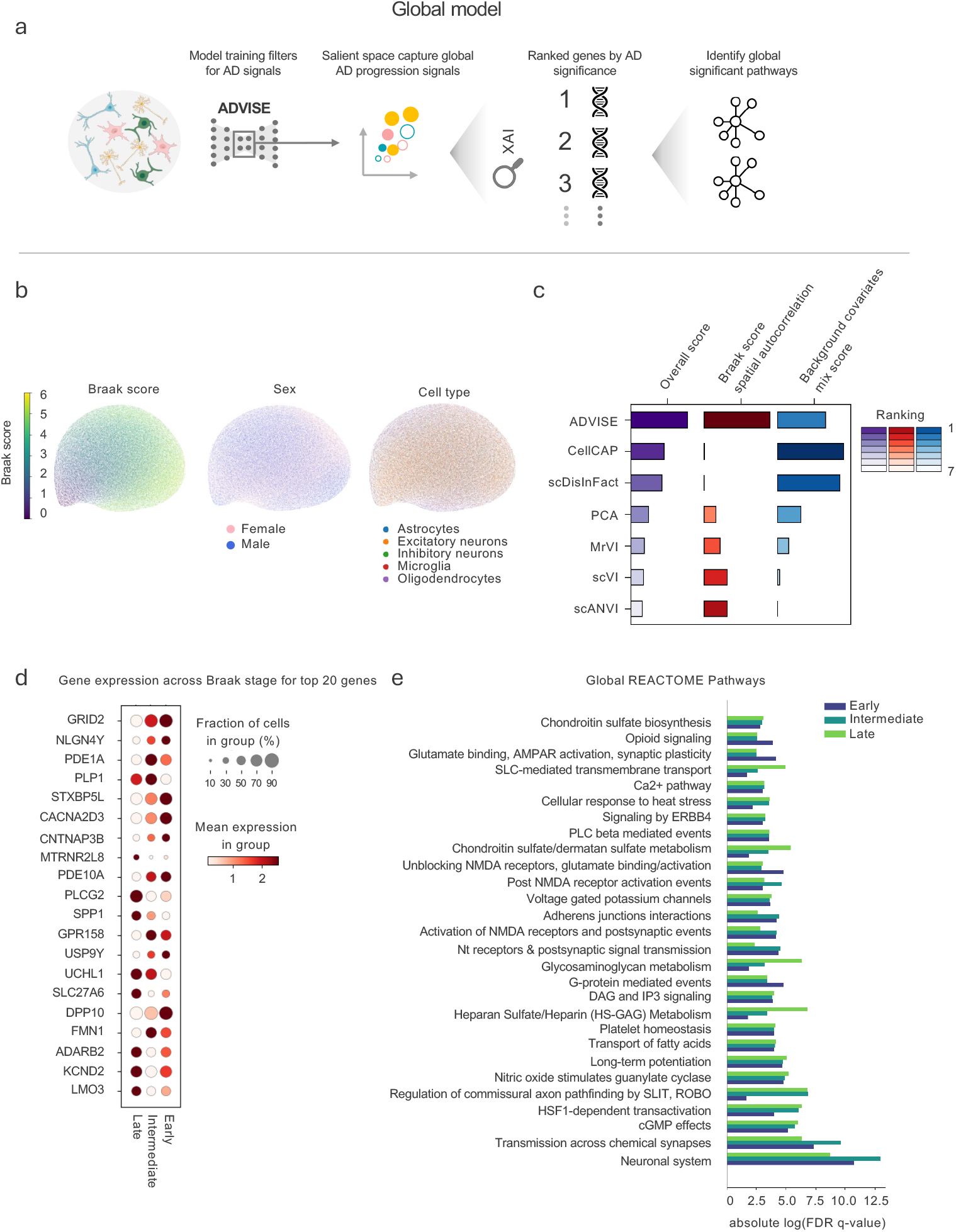
ADVISE captures global AD progression signatures. **(a)** Overview of the global ADVISE workflow. The ADVISE model is trained on all cells to isolate global AD-related signatures shared across sexes and cell types by treating cell type and sex as background covariates. The resulting AD variables capture gene expression signatures related to AD progression. Downstream XAI analyses applied to the AD variables identify global AD-related genes and pathways that are enriched throughout all sexes and cell types. **(b)** UMAP visualization of AD variables colored by Braak stage (left), sex (middle), and major cell types (right). **(c)** Benchmarking ADVISE against alternative dimensionality-reduction and integration methods (MrVI, scANVI, scVI, PCA, CellCap, scDisInFact) across multiple metrics, all scaled so higher is better. Each bar is a composite score, sized by its relative value and colored by the method’s ranking. Braak Stage Spatial Autocorrelation averages Geary’s C and Moran’s I; Background Traits Mix Score averages cell-type and sex entropy of mixing; Overall Score averages the two. **(d)** Normalized expression patterns for the top 20 AD-associated genes identified by ADVISE. The gene expression patterns are stratified across Braak stages. The size of each dot represents the fraction of cells expressing the gene, and the intensity of the color represents the mean expression value. **(e)** Significantly enriched AD-related pathways involved in key biological processes of AD progression, based on the REACTOME database. Pathway enrichment analysis was performed on ADVISE ranked list of global AD genes to highlight top pathways that are involved in 6 AD progression. Significant pathways were selected by average significance across Braak stages (Benjamini-Hochberg FDR q *<* 0.05).

We next compared ADVISE to several existing baseline methods, including MrVI [30], scANVI [21], CellCap [31], and scDisInFact [32], which incorporate Braak score labels during training, as well as scVI [20] and principal component analysis (PCA), which are unsupervised single-cell representation learning approaches. Quantitative evaluation demonstrated that ADVISE outperforms all of the baseline methods across multiple metrics, including spatial autocorrelation (Methods; Geary’s C and Moran’s I) between learned AD embeddings and Braak score, as well as mixing scores (Methods; Entropy of mixing) across covariates including sex and cell type (Figure 2c). Together, these results provide confidence that ADVISE consistently isolates Braak score-associated signals from confounding background variation.

To interpret the molecular drivers of AD progression captured by the model, we applied Expected Gradient (EG) [33] to identify genes that contribute most strongly to changes along the disease trajectory. We then performed gene set enrichment analysis (GSEA) using REACTOME pathways on the ADVISE-ranked gene list to characterize the dominant biological processes associated with different stages of AD (Methods; Pathway analysis). Among the top 20 genes identified by the model (Figure 2d), we observed two Y-chromosome genes: *USP9Y* and *NLGN4Y. USP9Y* has been implicated in Tau phosphorylation and may contribute to sex-specific molecular differences in Tau pathology [34], while *NLGN4Y* encodes a neuroligin family protein involved in postsynaptic cell adhesion and has been linked to cognitive and neurodevelopmental disorders [35]. The identification of Y chromosome-linked genes is consistent with known sexual dimorphism in AD progression [36]. In addition, a substantial fraction of the top-ranked genes are involved in synaptic function and signal transduction, including *PLP1* and *PDE1A. PLP1* encodes the major protein component of myelin, and its aberrant expression has been associated with increased neuroinflammation and elevated AD risk [37]. *PDE1A* encodes a phosphodiesterase that regulates intracellular cAMP levels, and dysregulated cAMP signaling has been linked to multiple neurodegenerative disorders, including AD [38].

We then identified multiple significantly enriched pathways (Figure 2e). In agreement with the top gene-level findings, many of the most enriched pathways are associated with synaptic and neuronal function, including Unblocking of NMDA Receptors, Voltage-Gated Potassium Channels, Long-Term Potentiation, Transmission Across Chemical Synapses, and the Neuronal System. We also identified enrichment of the HSF1-Dependent Transactivation pathway, which is involved in cellular stress responses, highlighting the impact of increased Tau burden on neuronal homeostasis. Additionally, the Transport of Fatty Acids pathway emerged as significantly enriched, which is consistent with known disruptions to cellular metabolism and bioenergetics during AD progression [39, 40]. The pathway is also implicated in axonal myelination, which is known to be dysregulated in AD [41]. The convergence of biologically relevant genes and pathways provides further evidence that ADVISE captures meaningful molecular signatures underlying AD progression.

### ADVISE e!ectively captures AD signals associated with sex

While the global model captures AD signals shared across contexts, substantial evidence indicates that AD pathology differs by sex [18, 42, 11]. Sex-specific molecular signatures may be diluted when disease mechanisms are analyzed at a global scale. To recover these sex-specific signatures, we constructed a sex-specific model (Figure 3a) to resolve differences in AD progression between males and females. In this model, ADVISE was trained on the unstratified dataset, with Braak score and sex encoded in the AD latent space and cell type assigned to the background space. In doing so, the latent embeddings are separated by sex within the AD variable space, while preserving a clear trajectory of Braak stage progression within each group (Figure 3b).

**Figure 3.**
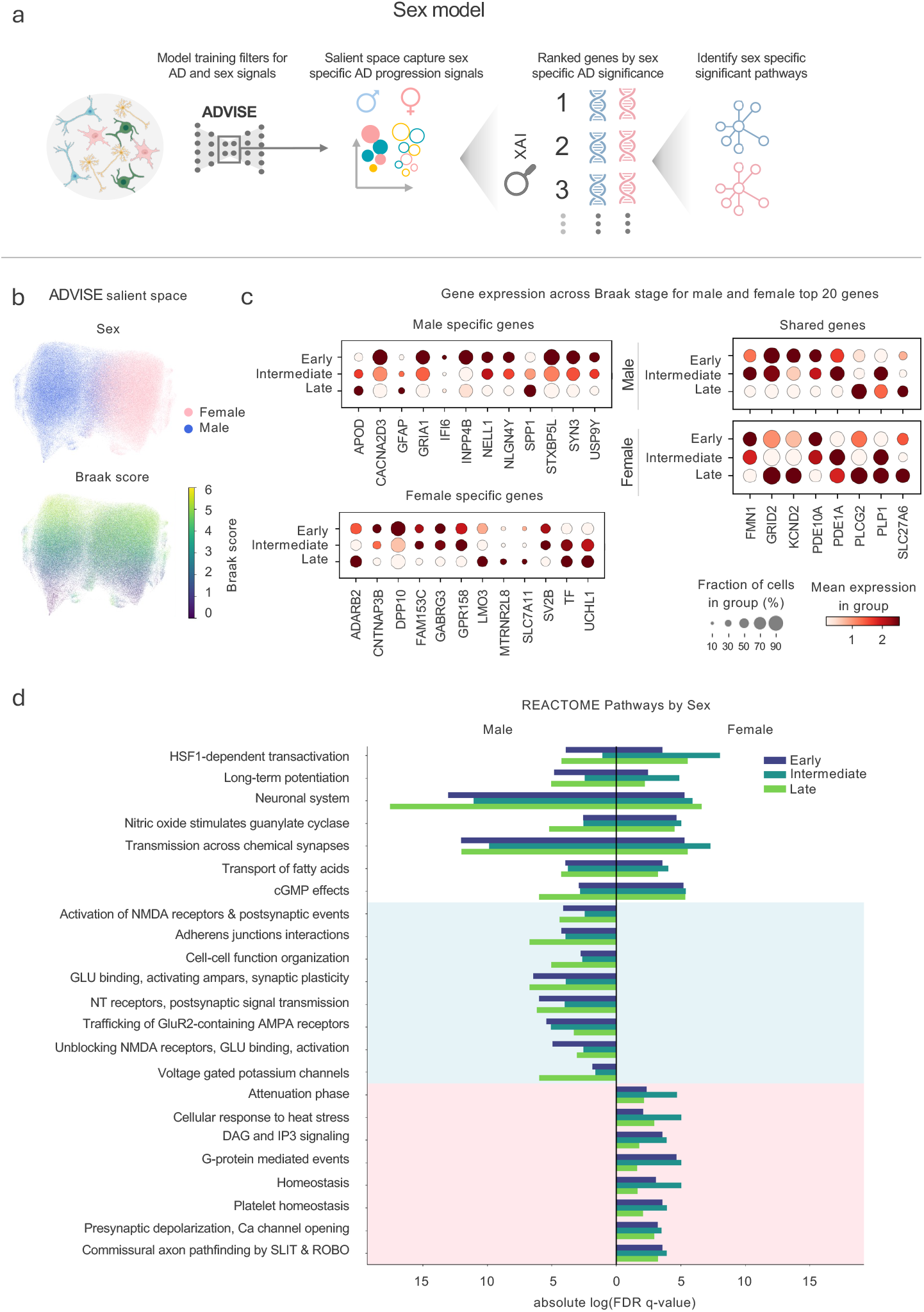
ADVISE identifies sex-specific AD progression signatures. **(a)** Overview of the sex-specific ADVISE modeling workflow. The ADVISE model is trained on all cells to isolate sex-specific signatures by predicting Braak score and sex from the AD variables, while treating cell type as the background covariate. The resulting AD variables capture gene expression signatures according to sex-specific contributions to AD progression. Downstream XAI analyses are applied separately to male and female cells using the AD variables, enabling the identification of genes and pathways with sex-specific AD-related dysregulation. **(b)** UMAP visualization of ADVISE AD variables colored by sex and Braak stage. **(c)** Normalized expression patterns of the top 20 AD-associated genes identified by ADVISE. **Left:** Sex-specific genes among the top 20 for each sex. **Right:** Genes shared between the top 20 lists of both sexes. The gene expression patterns are stratified by Braak stage. Dot size indicates the fraction of cells expressing the gene, and color intensity represents mean expression. **(d)** Significantly enriched AD-related pathways involved in key biological processes of AD progression, grouped into common, male-specific, and female-specific pathways based on the REACTOME database. Pathway enrichment analysis was performed on ADVISE ranked list of AD genes for each sex to highlight pathways that are significantly involved in AD progression. Significant pathways are chosen based on the average significance across each Braak stage. Significance was assessed at 8 FDR q *<* 0.05 using the Benjamini-Hochberg correction.

Using the same explainability pipeline as the global model, we ranked genes by their contribution to disease progression in each sex. Comparison of the top ranked genes revealed both shared and sex-specific signatures, motivating a focused analysis of sexually dimorphic genes (Figure 3c). Among genes shared across both sexes, *PLP1* and *PDE1A* again emerged as highly ranked, consistent with the global model findings (Figure 3c, right). *FMN1* was also highlighted in both sexes. Notably, supplementation of *FMN1* in yeast mitigates A*ω* toxicity, suggesting a protective role in AD progression [43].

Male specific gene signatures included *APOD*, which is elevated in AD patients and harbors polymorphisms associated with increased sporadic AD risk in a sex dependent manner [44]. The *APOD* promoter contains estrogen, glucocorticoid, and progesterone response elements, suggesting hormone mediated regulation that differs between males and females [45]. *SPP1*, another male specific gene highly ranked by the model, has been linked to faster cognitive decline and increased neuropathological burden [46]. Its expression is suppressed by estrogen exposure in mice, providing a potential mechanism for sex-specific regulation [47]. In addition, Y chromosome genes *USP9Y* and *NLGN4Y* were identified, consistent with sex-linked effects also observed in the global model (Figure 3c, left).

Female specific genes shared common themes of epigenetic and hormone-sensitive regulation. The methylation status of *ADARB2* has been associated with sex-specific AD pathology, with increased methylation observed in AD positive males but not in females [48]. *GPR158*, a G protein coupled receptor involved in synaptic modulation, has been implicated in AD and is downregulated by estrogen antagonists in breast cancer, suggesting estrogen dependent regulation in the brain [49]. *MTRNR2L8*, a predicted negative regulator of apoptosis, displays sexually dimorphic expression in AD patients [18]. Finally, *SV2B*, a binding partner of *BACE1* that suppresses A*ω* production, exhibits sexually dimorphic expression in human tissues, raising the possibility of similar regulation in the brain [50, 51] (Figure 3c, left).

We next examined biological pathways enriched by sex (Figure 3d). Pathways shared across both sexes largely recapitulate those identified in the global model, including cellular stress responses such as HSF1 Transactivation and metabolic processes such as Fatty Acid Transport. Long Term Potentiation was also enriched in both sexes, consistent with extensive evidence of synaptic plasticity deficits in AD [52].

Male specific pathways were dominated by synaptic and electrophysiological processes. NMDA Receptor Activation and downstream Postsynaptic Signaling are central to neural plasticity and have been strongly implicated in AD pathogenesis [53]. Emerging evidence suggests that kinases downstream of NMDA receptors are regulated in a sex dependent manner [54]. Alterations in glutamatergic signaling, including trafficking and activation of *GluR2* containing *AMPA* receptors, are also well documented in AD, with disease associated downregulation of *GluR2* [55]. In rodents, *GluR2* has been shown to have decreased expression in male rats compared to female rats, suggesting a sex-specific mechanism regulating *GluR2* expression that may be conserved in humans [56]. In addition, Voltage-gated Potassium Channels, which regulate neuronal excitability and survival, have been linked to AD related apoptosis. Their function is regulated in a sex dependent manner under stress, with males showing greater potassium leakage, suggesting heightened susceptibility in males [57, 58].

Female specific pathways were enriched for hormone-sensitive signaling related to axon guidance and calcium homeostasis. *SLIT/ROBO* Mediated Commissural Axon Pathfinding is of particular interest, as *APP* functions as a *SLIT* receptor in *C. elegans*, raising the possibility that altered *APP* processing in AD disrupts this pathway [59]. Expression of *SLIT* and *ROBO* genes is hormonally regulated in human reproductive tissues, suggesting analogous regulation in the brain [60]. In parallel, *DAG* and *IP3* signaling, which governs intracellular calcium release and downstream nuclear signaling, has long been implicated in AD through dysregulation of calcium homeostasis [61]. This pathway is modulated by estrogen exposure, reinforcing its sexually dimorphic nature [62]. Consistent with this, Presynaptic Depolarization and Calcium Channel Opening, key steps in neurotransmitter release, are also sensitive to estrogen mediated regulation, highlighting a female specific vulnerability rooted in hormone regulated calcium signaling [62].

### ADVISE enables comparison of cell type-specific AD signals

Many transcriptional signatures associated with AD are highly cell type-specific [7, 15, 63, 64]. As a result, models trained across heterogeneous cellular populations can obscure disease relevant signals that manifest differently across cell types. To address this, we trained cell type-specific ADVISE models that explicitly filter for AD signatures significant within each individual cell type (Figure 4a). In this framework, ADVISE trained separate, cell type-specific models on the stratified dataset with Braak score encoded in the AD latent space and sex assigned to the background space (Methods; Model optimization details). The cell type-specific ADVISE models produced UMAP embeddings with smooth, continuous Braak score trajectories within each cell type (Figure 4b). These structured gradients reflect a coherent progression of disease severity and indicate that ADVISE successfully captures AD-related transcriptional signals without conflating them across diverse cellular populations.

**Figure 4.**
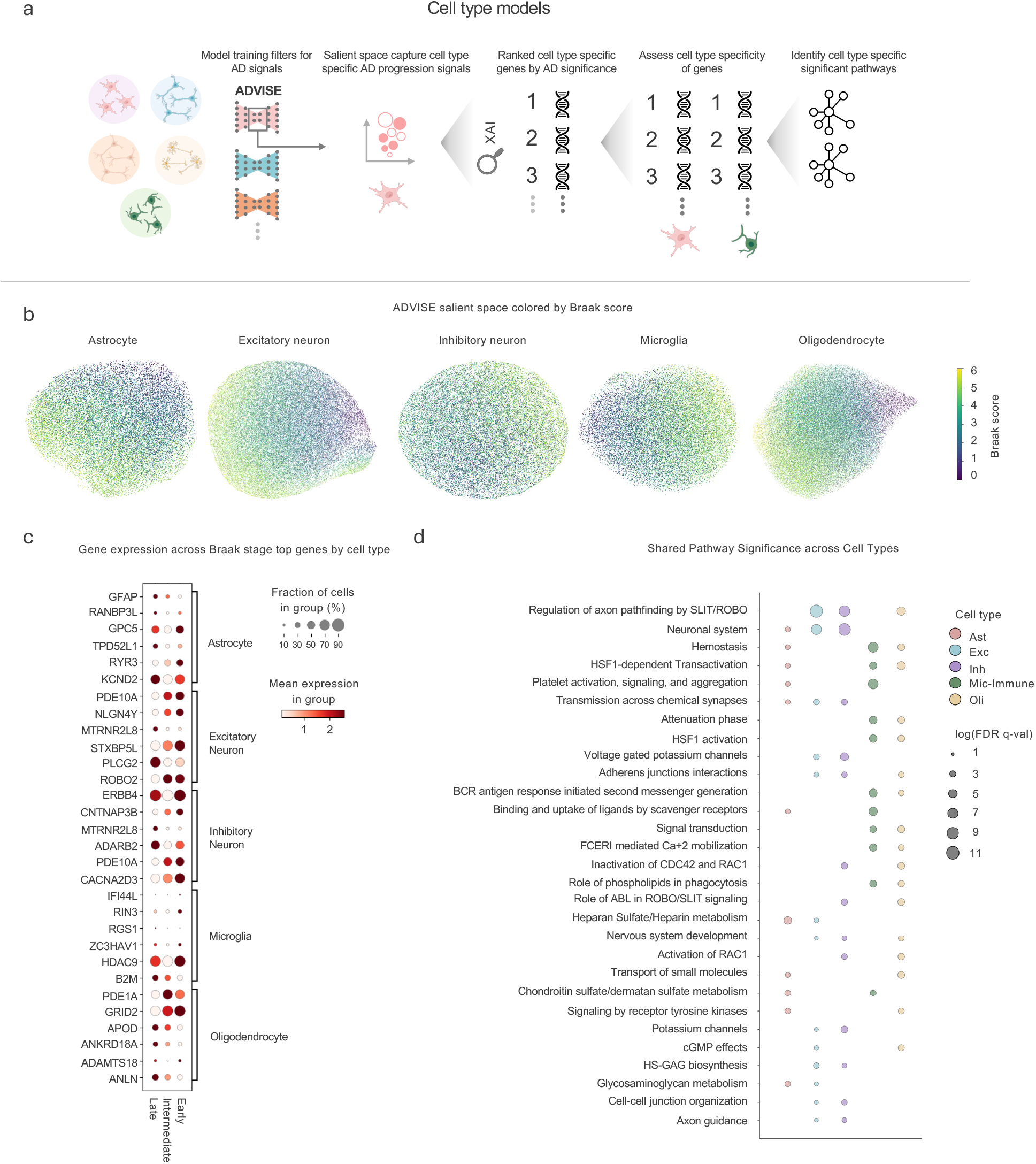
ADVISE identifies cell type-specific AD progression signatures. **(a)** Overview of the cell type-specific ADVISE modeling workflow. Separate ADVISE models are trained on cells from each cell type to isolate cell type-specific AD signatures by predicting Braak score from the AD variables, while treating sex as the background covariate. The resulting cell type-specific AD variables are subsequently analyzed using downstream XAI methods to identify AD-related genes and pathways specific to each cell type. **(b)** UMAP visualization of ADVISE AD variables for each major cell type colored by Braak stage. **(c)** Normalized expression patterns of selected AD-associated genes identified by ADVISE. Identified genes were chosen from the top 20 list of ADVISE ranked AD genes for each cell type. Gene expression is stratified by cell type and Braak stage. Dot size indicates the fraction of cells expressing the gene while color intensity represents mean expression. **(d)** Dot plot summarizing pathways that are significantly enriched across multiple cell types. Dot size indicates absolute ***−****log*_10_(FDR q-valu10e) significance of each pathway, and color denotes the cell type in which the pathway becomes enriched. Significant pathways are chosen based on the average significance across each Braak stage. Significance was assessed at FDR q *<* 0.05 using the Benjamini-Hochberg correction.

We first identified cell type-specific genes that contributed to AD progression (Figure 4c). Several genes previously detected in the global and/or sex-specific analyses were also recovered here, demonstrating consistency of key AD-associated signals across analytical scales. These overlapping genes include *NLGN4Y* in excitatory neurons, *PDE1A* and *APOD* in oligodendrocytes, and the potassium channel *KCND2* in astrocytes. Their recurrence across modeling strategies suggests that they represent robust transcriptional features of AD that persist even after explicitly accounting for cellular heterogeneity. Beyond these shared signals, distinct neuronal patterns emerged in the gene findings. Both excitatory and inhibitory neurons exhibited multiple significant genes related to synaptic organization and signaling, underscoring the central role of synaptic dysfunction in AD. In excitatory neurons, *ROBO2* was particularly notable, directly aligning with the enrichment of the SLIT/ROBO axon guidance pathway observed in the cell type-specific pathway analysis (Figure 4d). This concordance between gene- and pathway-level results highlights disrupted axon guidance and neuronal connectivity as key features of AD progression in excitatory populations. In inhibitory neurons, prominent synapse-associated genes included *ERBB4*, a tyrosine kinase receptor involved in neuronal signaling and synaptic modulation. Given prior evidence linking *ERBB4* to AD development [65], its identification here further supports a role for altered inhibitory synaptic signaling in disease progression.

Glial cell types showed patterns consistent with immune and structural regulation. In microglia, one of the most significant genes was *HDAC9*, which encodes a histone deacetylase belonging to a family of epigenetic regulators widely implicated in AD [66]. Notably, downregulation of *HDAC9* has been associated with neuroprotection in ischemic brain injury [67], suggesting a context-dependent role in neurodegenerative and neuroinflammatory processes. In astrocytes, *GFAP*, a cytoskeletal protein and established marker of astrocyte reactivity, was identified as significant. Elevated *GFAP* levels have been proposed as an early biomarker of AD [68], further linking our cell type-specific findings to clinically relevant disease signatures.

We then identified relevant biological pathways within each cell type and assessed their overlap to define cross cell type AD signatures (Figure 4d). The pathway with the highest combined significance across cell types was Axon Pathfinding Mediated by SLIT/ROBO genes, whose proteins are central to axon guidance, synaptic development, and broader processes such as cell proliferation and migration [69]. The prominence of this pathway highlights a conserved neurodevelopmental and synaptic component of AD pathology. Beyond this shared signal, we observed clear cell type-specific patterns. In both excitatory and inhibitory neurons, most significant pathways were related to synaptic function and neuronal signaling, including Voltage-gated Potassium Channels, Transmission across Chemical Synapses, the Neuronal System, and Adherens Junction Interactions. These findings underscore widespread disruptions to synaptic communication in neuronal populations during AD progression. In contrast, the dominant pathways identified in microglia were enriched for immune related processes, such as B Cell Receptor Antigen Response-initiated Second Messenger Generation and Platelet Activation and Aggregation, reflecting their specialized roles in neuroinflammation and immune surveillance in the AD brain.

Several of the enriched pathways were also identified in our prior global and sex-specific analyses, including Regulation of Axon Pathfinding by SLIT/ROBO, HSF1-Dependent Transactivation and HSF1 Activation, and cGMP-Related Signaling. Their recurrence across multiple scales reinforces the central role of these mechanisms in AD progression. The shared pathways also point to coordinated or complementary functions across cell types. For example, Regulation of Axon Pathfinding by SLIT/ROBO was significant in excitatory neurons, inhibitory neurons, and oligodendrocytes (cell types known to express ROBO/SLIT proteins), suggesting a broadly conserved disruption of axon guidance programs. Moreover, the BCR antigen response pathway, an immune-related pathway, was enriched in microglia and oligodendrocytes, consistent with their roles in immune regulation and inflammatory signaling. Excitatory neurons, inhibitory neurons, and oligo-dendrocytes all showed enrichment for Adherens Junction Interactions, a pathway critical to blood-brain barrier integrity, aligning with their established contributions to barrier structure and maintenance.

We next examined transcriptional features uniquely enriched within each cell type. We compared AD-associated signatures across cell types and prioritized pathways and genes with strong cell type-specific effects (Figure 5). We selected pathways that were not only statistically enriched but also closely aligned with the established biological roles of the corresponding cell type, allowing us to interpret how AD related dysregulation manifests in distinct cellular contexts. In parallel, we quantified gene level specificity by measuring, for each gene, the difference in AD-associated significance between a given cell type and the next most significant cell type. Genes were ranked by this differential score, and the top candidates were defined as the most cell type-specific. Together, this approach isolates uniquely perturbed molecular programs that may be masked when focusing solely on shared AD signatures.

**Figure 5.**
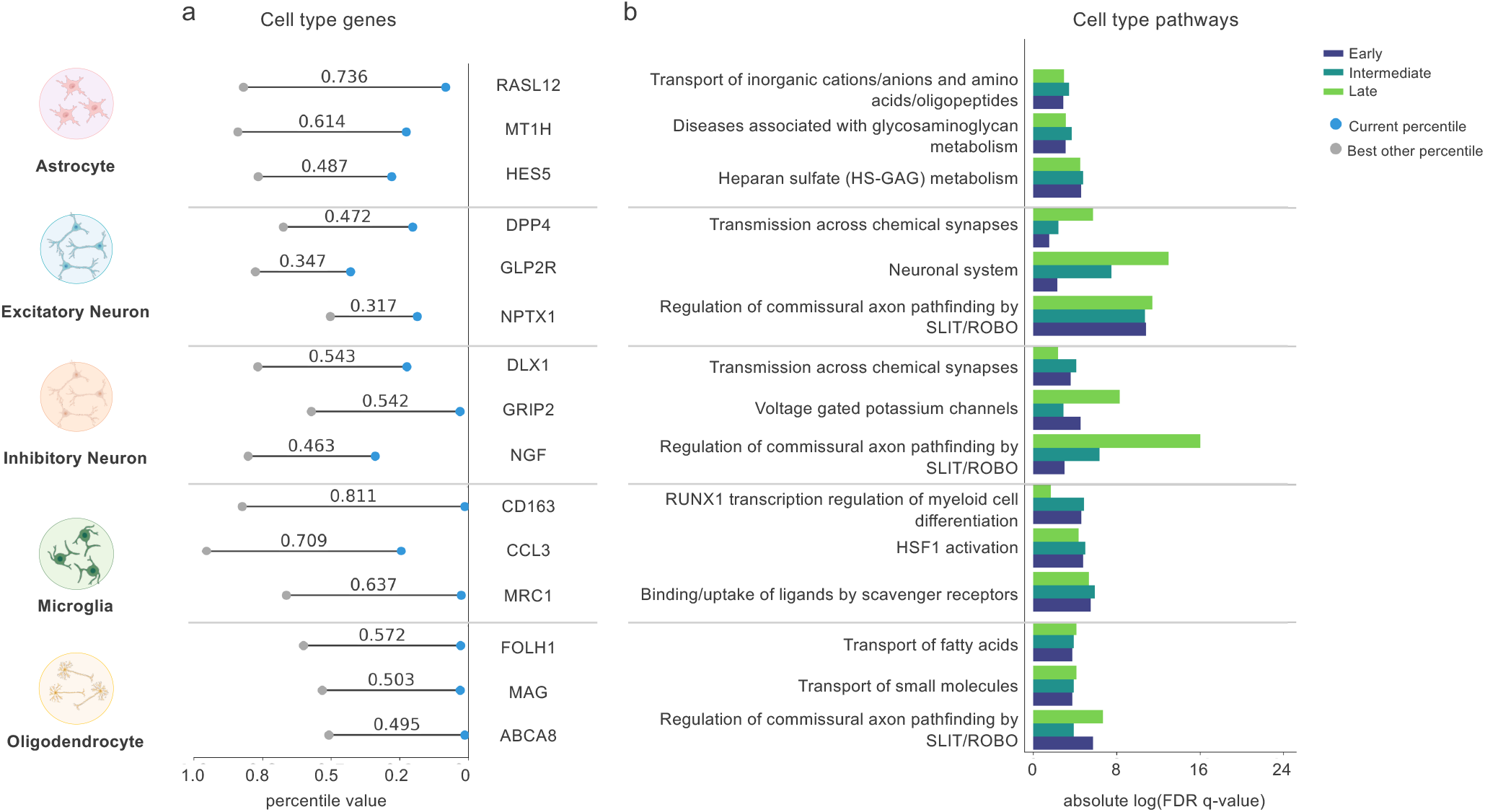
Cell type analysis of gene specificity and top contributing pathways to AD progression. **(a)** Top cell type-specific genes related to AD progression identified by the models. For each gene, the percentile value within the original cell type (blue marker) is shown alongside the highest percentile observed across all other cell types (gray marker), enabling quantification of enrichment differences across cell types. Identified genes were selected from a list of ADVISE ranked AD genes for each cell type. The top three genes with the highest percentile difference were highlighted for each cell type. **(b)** Top cell type-specific REACTOME pathways that are significantly enriched and involved in key biological processes related to AD progression. Significant pathways are selected based on the average significance across Braak stages. Bar length indicates the absolute significance value, and bar color denotes the Braak stage. Significance was assessed at FDR q *<* 0.05 using the Benjamini-Hochberg correction.

Astrocyte specific pathways were tightly aligned with their core roles in maintaining synaptic and extra-cellular homeostasis. Enrichment of Transport of Inorganic Cations, Anions, and Amino Acids is consistent with astrocytic functions in potassium buffering and glutamate clearance processes known to be disrupted in AD and linked to neuronal hyperexcitability. Pathways related to Heparan Sulfate/Heparin Glycosaminoglycan (HS-GAG) metabolism further point to altered extracellular matrix regulation. Astrocytic heparan sulfate proteoglycans influence growth factor signaling, synaptic stability, and the aggregation and spread of amyloid-*ω* and Tau, suggesting that extracellular remodeling is a key astrocyte-specific component of AD pathology. The top astrocyte specific genes reinforce these functional themes. *MT1H*, a metal binding protein, aligns with disrupted ionic regulation and has been reported to be elevated in AD [70]. *HES5*, a regulator of NOTCH signaling, connects to established AD related signaling changes and is consistent with interactions between *NOTCH* and heparan sulfate pathways [71]. *RASL12*, a marker of toxic astrocytes associated with disease severity [72], further supports a link between astrocyte reactivity and AD progression.

In contrast, neuronal populations exhibited predominantly synaptic and connectivity-related signatures. Both excitatory and inhibitory neurons were enriched for pathways involving Transmission across Chemical Synapses and SLIT/ROBO axon pathfinding. The repeated identification of the SLIT/ROBO axon pathfinding pathway, also observed in global and sex-specific analyses, highlights disrupted neuronal connectivity as a robust and multi-scale feature of AD. Its enrichment in both neuronal and oligodendrocyte populations further suggests coordinated dysregulation of axon guidance and circuit integrity. Gene-level specificity refined these neuronal distinctions. In excitatory neurons, *DPP4*, a therapeutic target under investigation for AD [73], and *NPTX1*, observed near amyloid plaques and implicated in amyloid-*ω*-mediated synaptic effects [74], were among the top candidates. In inhibitory neurons, *DLX1*, involved in interneuron differentiation and reported to be upregulated in AD [75], and *GRIP2*, shown to be downregulated in AD neurons [76], emerged as specific signals. Impairment of NGF-related signaling in AD [77] further supports selective vulnerability of inhibitory circuits.

Microglia specific pathways were predominantly those involved in immune function regulation. Binding and Uptake of Ligands by Scavenger Receptors, HSF1 Activation, and *RUNX1* Regulates Transcription of Genes involved in differentiation of myeloid cells are all pathways involved in pathogen recognition, cell stress response, and inflammation. Immune dysregulation in the brain is a key hallmark of AD pathology which is corroborated by these results. Microglia specific gene analysis further expanded on the immune function signature from the pathway analysis. *CD163*, a scavenger receptor protein, has been associated with small vessel injuries in mild cognitive impairment and has shown increased expression in AD [78]. *CCL3* is a proinflammatory cytokine that has been shown to play a role in neuroinflammation in mice [79]. *MRC1* is a mannose receptor involved in pathogen identification and engulfment that is upregulated in mouse models of AD [80].

Oligodendrocyte-specific pathways were indicative of their role as neuronal support cells. Enrichment of Transport of Fatty Acids and Regulation of Commissural Axon Pathfinding by *SLIT* and *ROBO* highlight this by identifying changes in neuron myelination and neuronal synaptic maintenance in AD, respectively. Oligodendrocyte-specific gene analysis complements the pathway analysis by identifying several genes involved in neuronal myelination maintenance, such as *ABCA8* and *MAG*, where the process of neuron myelination is known to be disrupted in AD [81]. Consistent with neuronal maintenance is the identification of the *FOLH1* gene, which is involved in glutamate metabolism and has been identified as an early onset risk gene for AD [82].

### Experimental validation of *ROBO2*

In our cell type-specific analysis, we observed a trend of decreased *ROBO2* expression in excitatory neurons at late Braak stages (Figure 4c). Additionally, *ROBO/SLIT* signaling pathways were identified as significant across several different analyses (Figure 2e, 3d, 5b). This pathway has been previously implicated in AD pathogenesis [83], although its role in relation to Tau pathology is not well described. Because our data suggested a Braak-stage-dependent down-regulation of *ROBO2*, we tested whether activation of this pathway functionally impacts Tau in human excitatory cortical neurons derived from hiPSCs (Figure 6a). To test this, we engineered a hiPSC line that expresses a doxycycline-inducible dCas9-VPR (CRISPR activation, CRISPRa). hiPSCs were transduced with a lentiviral construct containing a gRNA targeting the *ROBO2* gene locus that was previously validated to efficiently upregulate *ROBO2* gene expression in HEK293T cells constitutively expressing dCas9-VPR. We differentiated these hiPSCs to excitatory neurons following established protocols [84, 85] and documented increased ROBO2 protein abundance in hiPSC-derived neurons by western blot and immunocytochemistry (Supplementary Figure 4a-c). To assess effects on Tau abundance and phosphorylation in these neurons we analyzed phospho-Thr 231 levels as well as total Tau levels in neurons with *ROBO2* upregulation compared to a non-induced control. In neurons with upregulated *ROBO2*, we observed an increase in total Tau but no change in p-Tau Thr 231, resulting in an overall decrease in the p-Tau Thr 231/total Tau ratio (Figure 6b-d). Hyper-phosphorylation of Tau is a hallmark pathology of AD, thus activating *ROBO2* may represent a protective mechanism.

**Figure 6.**
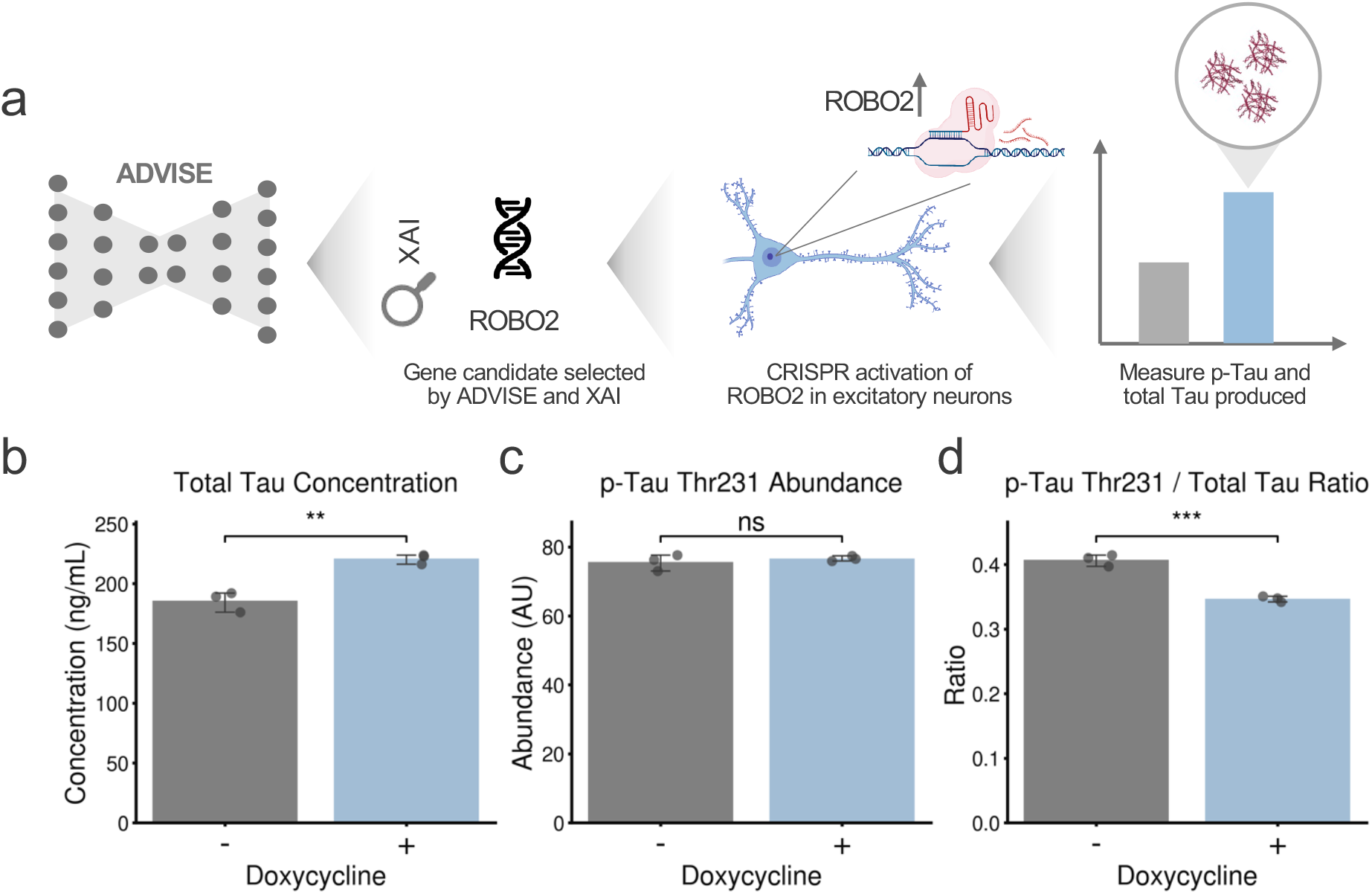
Impact of *ROBO2* upregulation on total Tau and pTau Thr 231 abundances for hiPSC-derived neurons. **(a)** Overview of experimental validation pipeline. The *ROBO2* gene candidate was chosen from a list of ADVISE-identified top genes. CRISPR activation was then performed to increase *ROBO2* expression in hiPSC-derived neurons, and resulting p-Tau and total Tau produced were measured against a control. **(b)** Total Tau, **(c)** p-Tau Thr 231, and **(d)** p-Tau Thr 231/Total Tau ratio for hiPSC-derived neurons treated with doxycycline compared to a non-treated control. Individual data points represent expression values from independent wells. ** P *<* 0.01, *** P *<* 0.001

## Discussion

ADVISE identified coordinated changes in synaptic transmission, neuronal excitability, HSF1-dependent stress responses, and metabolic regulation as defining features of AD progression, resolving these programs across global, sex-specific, and cell-type-specific contexts. At the global level, the prominence of synaptic and bioenergetic pathways aligns with established models of synaptic failure and metabolic stress in AD. Critically, these signals became detectable only after Braak stage-associated variation was disentangled from dominant covariates such as sex and cell type, demonstrating that unmodeled biological confounders can obscure disease-relevant signals in single-cell data. Across all scales, synaptic and stress-related programs followed non-monotonic trajectories over Braak stage rather than changing linearly, pointing to stage-specific compensatory or maladaptive responses during disease progression. Most notably, ADVISE repeatedly prioritized the SLIT/ROBO axon guidance pathway and nominated *ROBO2* as a progression-associated candidate in excitatory neurons. Experimental validation demonstrated that increasing *ROBO2* expression in excitatory neurons decreased the ratio of phosphorylated to total Tau, supporting a protective role against AD progression and nominating a novel therapeutic gene target.

Compared with the original findings of Mathys et al., our results show partial concordance between the two approaches, with stronger agreement observed in glial populations and weaker correspondence in neuronal cell types (Supplementary Figure 3). This trend is reflected both in rank correlations and in the limited overlap among top-ranked genes, which remains consistently modest across all cell types. Together, these patterns suggest that ADVISE does not simply reproduce previously reported transcriptional programs, but instead yields a partially overlapping yet distinct prioritization of genes, with the greatest divergence emerging in neuronal compartments.

Several limitations should be considered in interpreting these results within the context of AD. First, ADVISE is inherently data-driven and relies on the quality, scale, and annotation of available single-cell datasets. While the Mathys et al. atlas provides unprecedented coverage, certain axes of variation remain underrepresented. For example, attempts to disentangle aging specific signals from AD-specific pathology were constrained by a limited and uneven age distribution, preventing robust modeling of normative aging trajectories independent of disease. In addition, Braak staging reflects Tau pathology but does not fully capture amyloid burden, vascular contributions, or cognitive status, potentially limiting the scope of inferred progression signals. In terms of experimental validation, the hiPSC-derived neurons display a fetal-like transcription profile. In the context of AD, an aging-associated disease, this model may miss aspects of disease impacted by transcriptional changes due to aging. Additionally, we tested *ROBO2* gene upregulation in a single-cell (excitatory neuron) system. It is reasonable to suggest that the effect of upregulating this gene may be altered in a multicellular context.

An important future extension of ADVISE is to disentangle transcriptional programs associated with neuropathology from those linked to cognitive resilience. A subset of individuals exhibit high AD pathology yet minimal cognitive decline [86, 87, 88]. With sufficiently representative datasets, ADVISE could be applied to compare these resilient individuals with cognitively impaired patients at similar Braak stages, potentially revealing protective gene expression programs and novel therapeutic strategies focused not only on pathology reduction but also on preservation of cognitive function.

ADVISE provides a scalable framework for disentangling complex sources of biological variation in single-cell datasets to reveal transcriptional programs associated with Alzheimer’s disease progression. By explicitly modeling and separating Braak stage effects from dominant confounders such as sex and cell type, the model enables identification of disease-associated signatures that are otherwise obscured in standard analyses. Applying this framework uncovered multiscale patterns of AD progression, including global stress and synaptic dysfunction programs, sex-specific pathway vulnerabilities, and cell type–resolved regulatory signatures. Notably, ADVISE highlighted recurrent dysregulation of the SLIT/ROBO axon guidance axis, with *ROBO2* emerging across multiple contexts and validated experimentally in excitatory neurons. Beyond these specific findings, the model establishes a general strategy for integrating heterogeneous single-cell data to identify stage-resolved and context-specific molecular mechanisms of neurodegeneration.

## Methods

### The ADVISE model

Here, we present the ADVISE model in more detail. The model’s generative process extends from the ACE base model [22].

#### The ADVISE generative process

For a data point *x*_*n*_, we assume that each expression value *x*_*ng*_ for sample *n* and gene *g* is generated through the following process:

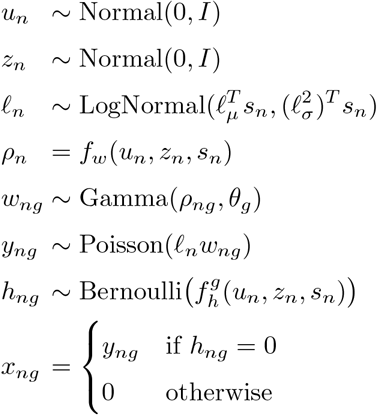

We introduce two latent variables, *u*_*n*_ and *z*_*n*_, to capture variation in scRNA-seq data. The variable *u*_*n*_ models variation associated with Braak score (*a*_*n*_), isolating AD-related biological signals, while *z*_*n*_ captures background variation driven by covariates *b*_*n*_ (e.g., cell type, sex), which often dominate and obscure subtle disease signals. This design enables disentanglement of AD-related and background effects across complex biological contexts. Both *u*_*n*_ and *z*_*n*_ are assigned standard multivariate Gaussian priors to facilitate efficient inference within the VAE framework [89].

To enforce that *u*_*n*_ and *z*_*n*_ capture distinct sources of variation, we use two complementary strategies. First, we introduce supervised prediction losses: a network predicts Braak score (*a*_*n*_) from *u*_*n*_, and another predicts background covariates *b*_*n*_ from *z*_*n*_. Depending on the task, we use cross-entropy (classification) or mean squared error (regression). Second, we apply the *d*-variable Hilbert-Schmidt Independence Criterion (*d*HSIC) to enforce statistical independence between *u*_*n*_ and *z*_*n*_. Minimizing *d*HSIC ensures the latent spaces encode non-overlapping biological signals, effectively disentangling AD-related variation from background effects.

We model library size using a batch-specific log-scale prior parameterized by *l*_*µ*_, 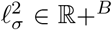, where *B* is the number of batches and *s*_*n*_ is a one-hot batch indicator. These parameters are set to the empirical mean and variance of log library size per batch. Gene expression is modeled using a Gamma distribution with mean *ρ*_*ng*_ and gene-specific shape parameter *θ*_*g*_, which can be interpreted as an inverse dispersion parameter of a negative binomial distribution and is learned via variational inference. Neural networks *f*_*w*_ and *f*_*g*_ map latent variables and batch annotations to gene space (*f*_*w*_: ℝ^*d*^ × {0, 1}^*B*^ → ℝ^*G*^ and similarly for *f*_*g*_, where *d* is the combined dimensionality of the concatenated salient and background latent spaces). The output of *f*_*w*_ represents gene expression proportions and is constrained via a softmax so that 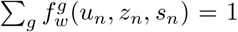. A separate network, *f*_*h*_, models dropout events due to technical noise.

Our generative process builds on contrastiveVI [23], with the addition of prediction networks and a *d*HSIC penalty. Unlike contrastiveVI, which requires explicit case-control groups, ADVISE leverages continuous covariates (*a*_*n*_, *b*_*n*_) to isolate Braak-related variation without requiring matched control cells.

### Datasets and preprocessing

To evaluate Braak stage prediction in the human brain, we used a prefrontal cortex single-nucleus RNA-sequencing dataset [7]. Following the original preprocessing pipeline, we removed droplets and filtered out cells with fewer than 200 detected genes. To ensure sufficient representation across cell populations, we retained the five most abundant cell types. We then identified the top 2,000 highly variable genes using the Seurat v3 method, applying batch correction via the batch key to raw counts. Because mitochondrial transcripts are indicative of poor single-nucleus sequencing quality, these genes were excluded. Finally, we randomly held out 20% of cells as a test set to enable unbiased evaluation.

For external validation, we used an independent single-nucleus RNA-sequencing dataset from an Alzheimer’s disease cohort that was not seen during model training [16]. This dataset was preprocessed using the same pipeline described above. After preprocessing, it comprised approximately 1.2 million cells from 84 patients. We then trained a global model using the same procedure and performed EG analysis to obtain ranked gene lists. To assess consistency between datasets, we computed the Spearman correlation between the ranks of overlapping genes from the external and original global models, yielding a high correlation of 0.87 (Supplementary Fig. 2).

### Model optimization details

For all datasets, ADVISE models were trained on 70% of cells, with 10% used for validation to determine the optimal number of training epochs, and the remaining 20% reserved for both quantitative and qualitative evaluation. Early stopping was applied when the validation ELBO failed to improve for 45 consecutive epochs. All models were optimized using the Adam optimizer with a learning rate of 0.0001, *ϵ* = 10^*−*8^, and a weight decay of 10^*−*5^. The hyperparameters λ_*a*_, λ_*b*_, λ_*d*HSIC_ were selected independently for each model to ensure that auxiliary loss terms were on a comparable scale to the ELBO.

Each ADVISE model was trained with 10 latent dimensions, with 3 allocated to AD-related variation and 7 to background variation, unless otherwise specified. For the global model, the target variable was Braak score, while cell type and sex were included as background covariates. In the sex-specific model, both sex and Braak score were treated as target variables, with cell type as the background covariate. For each cell type–specific model, Braak score served as the target variable and sex as the background covariate.

We compared ADVISE with the following baseline methods: MrVI [30], scANVI [21], scVI [20], CellCap [31], scDisInFact [32], and PCA. Each baseline was trained with 10 latent dimensions for consistency. For MrVI, Braak score was used as the sample key to capture disease progression effects. For scANVI, Braak score was treated as a categorical cell type label for conditional modeling. For CellCap, Braak score was used as the numeric target variable, with cell type and sex included as covariates. Similarly, for scDisInFact, Braak score was treated as the target condition, while sex and cell type were included as covariate conditions.

### Evaluation metrics

We evaluated model performance using quantitative metrics from the `scikit-learn` Python package. For visual comparison, we generated summary tables following contrastiveVI [23], displaying individual metrics and aggregate scores as bars. All metrics were min-max scaled to enable cross-metric comparison. We then computed two composite scores: *Braak Score Spatial Autocorrelation*, measuring the coherence of learned Braak-related structure, and *Background Covariates Mix Scores*, assessing how well cells from different background traits (e.g., cell type, sex) are mixed in the Braak latent space. The final overall score is the average of these two composites.

For *Braak Score Spatial Autocorrelation*, we computed both Geary’s C and Moran’s I to quantify spatial structure in the Braak-related latent variables learned by the global ADVISE model. Moran’s I captures global spatial trends, while Geary’s C is more sensitive to local differences between neighboring cells. The overall Braak score spatial autocorrelation is defined as the average of Geary’s C and Moran’s I.

For *Background Covariates Mix Scores*, we computed the Entropy of Mixing, which measures the diversity of background groups within each cell’s nearest neighbors in the learned Braak space. Higher entropy indicates better mixing of cells from different groups, suggesting effective removal of background-specific variation and improved isolation of Braak-related signals. We computed Entropy of Mixing for cell type and sex, and averaged these values to obtain the overall background mix score.

#### Geary’s C

To quantify local spatial autocorrelation in the learned Braak score latent space, we compute Geary’s C:

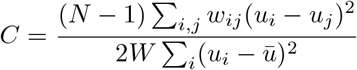

where *u*_*i*_ denotes the Braak score for cell *i, ū* is the global mean, *w*_*ij*_ is the spatial weight between cells *i* and *j*, and *W* = ∑_*i,j*_ *w*_*ij*_. We set *w*_*ij*_ = 1 if cell *j* is among the 50 nearest neighbors of cell *i*, and 0 otherwise. Because lower *C* values indicate stronger local autocorrelation, we report 1 *− C* so that higher values consistently reflect better performance.

#### Moran’s I

To assess global spatial autocorrelation in the Braak score latent space, we compute Moran’s I:

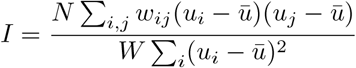

where *u*_*i*_ is the Braak score for cell *i, ū* is the global mean, and *w*_*ij*_ is defined as above. The denominator normalizes the spatial covariance of neighboring cells by the overall variance. Higher *I* values indicate stronger global spatial structure across the dataset.

#### Entropy of Mixing

To evaluate how well cells from different background covariates (e.g., cell types, sexes) are intermixed in the learned embedding space, we compute the entropy of mixing following prior work [20, 90]. For each cell *U*, we determine the empirical distribution of background groups among its 50 nearest neighbors, denoted *B*_*U*_. Let *p*_*i*_ be the fraction of neighbors belonging to group *i* out of *c* total groups, such that 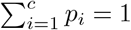. The local entropy of mixing for cell *U* is defined as:

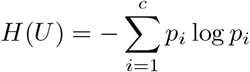

This calculation is repeated for 100 randomly selected cells, and the average entropy is reported. Higher values indicate better mixing of background groups, reflecting more effective removal of unwanted variation in the embedding.

### Interpretability of the ADVISE model

To interpret gene-level contributions to predicted Braak score, we applied the *Expected Gradients* (EG) method [33]. Expected Gradients generalizes Integrated Gradients by averaging over a distribution of background samples, yielding more stable and representative feature attributions for deep neural networks.

#### Definition

For a model *f* that maps an input gene expression vector *x* ∈ ℝ^*G*^ to a scalar output (here, predicted Braak score *â*), the Expected Gradient attribution for gene *g* is defined as:

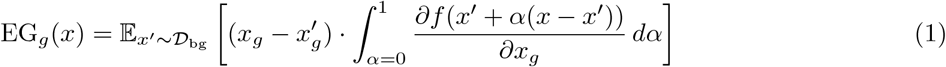

where: - *x* is a foreground (target) sample, - *x*′ is drawn from a background distribution *D*_bg_, - *g* indexes genes, - and the gradient is taken with respect to input gene *g*.

#### Foreground and Background Samples

In our framework, **Foreground samples** are 200 randomly selected cells, representing the cells whose predicted Braak scores we aim to interpret. **Background samples** are 800 randomly selected cells, forming the reference distribution *D*_bg_.

#### Application to Global, Cell Type, and Sex Models

For the *global model*, all cells were pooled regardless of cell type. EG values were computed using the aggregated foreground and background samples, and gene attributions were obtained by averaging the absolute EG values across all foreground cells.

For each *cell type model*, EG values were computed separately within each cell type. Foreground and background samples were drawn from the respective cell type-specific dataset, and genes were ranked by their mean attributions to reflect cell type-specific importance in Braak score prediction.

For the *sex model*, EG values were computed separately for each sex. Foreground and background samples were drawn from the respective sex-specific datasets, and genes were ranked by their mean attributions to reflect sex-specific contributions to Braak prediction.

### Pathway enrichment analysis

Pathway enrichment analysis is a computational approach used to assess whether a predefined gene set (i.e., a biological pathway) exhibits statistically significant expression differences between biological states. We performed this analysis using the open-source GSEAPY Python package with the prerank method. Pathways with a false discovery rate (FDR) q-value below 0.05, adjusted via the Benjamini-Hochberg procedure, were considered significantly enriched and are reported here.

The prerank method is a variant of Gene Set Enrichment Analysis (GSEA) that operates on a pre-ranked list of genes rather than raw expression values. In our study, genes were ranked according to their Expected Gradient values, reflecting their relevance to AD. GSEA then tests whether genes from a given pathway are disproportionately represented at the top or bottom of the ranked list, indicating enrichment beyond random expectation.

Enrichment analysis was conducted using the following gene set libraries:

- **KEGG (Kyoto Encyclopedia of Genes and Genomes):** A structured database of biological pathways covering metabolism, signal transduction, and disease mechanisms.
- **Reactome:** A manually curated knowledgebase describing molecular events in signal transduction, immune function, gene regulation, and other cellular processes.

While ranked gene lists can be generated per Braak stage, for ease of interpretation, we stratified Braak stages into early (0-2), intermediate (3-4), and late (5-6) stages. Gene attributions are generated across these stratified stages and then passed into pathway enrichment analysis to generate pathways per stratified disease stage.

### Comparison with original dataset paper

The original Mathys et al. study ranked differentially expressed genes between early- and late-pathology patients for each cell type based on statistical significance [7]. To compare these findings with those identified by our model, we evaluated both the overlap and the concordance of gene rankings for each cell type. Because our model outputs the top 2,000 genes per cell type, we restricted the Mathys gene lists to the top 2,000 genes to ensure a fair comparison.

#### Correlation analysis

For the subset of overlapping genes, we computed the Spearman rank correlation to assess the consistency between model-derived gene rankings and those reported in the original study.

### Wet lab validation

#### Cell culture

These experiments were performed using our well characterized human induced pluripotent stem cells (hiPSC) line [91] that was transduced with AAVS containing a doxycycline inducible dCas9-VPR by the University of Washington’s Ellison Stem Cell Core. hiPSCs were cultured in mTeSR plus medium and differentiated to neural progenitor cells (NPCs) following published protocols [92, 93] using dual SMAD inhibition via SB431542 (10 µM) and LDN-192189 (500 nM) treatment. Following 12 days of differentiation, NPCs were purified by sorting for CD24+/CD184+ cell populations as previously described [94]. NPCs were differentiated to neurons following our well characterized published protocol [95, 84, 85]. Briefly, NPCs were grown on 10 cm plates and were treated with GDNF (20 ng/mL), BDNF (20 ng/mL), and dbcAMP (250 µg/mL) for three weeks. Following the three-week differentiation, neurons were then treated with Accutase to dissociate into a single-cell suspension and replated on 24-well plates at a density of 5 × 105 cells per well. The following day, cells were treated with doxycycline (2 µg/mL) for 6 days to induce dCas9-VPR expression compared to vehicle control (DMSO) condition. For gRNA targeting validation experiments, HEK293T cells constitutively expressing dCas9-VPR were cultured in DMEM media containing 10% FBS. HEK293T cells were cultured in media containing puromycin (2 µg/mL) to select for cells that were transfected with gRNA-containing plasmids.

#### gRNA design and targeting validation in HEK293T cells

Five gRNA sequences targeting the *ROBO2* gene locus were designed using CRISPick software and Weissman lab resources for target validation in HEK293T cells. gRNA sequences tested in this study are shown in the below table. Forward and reverse gRNA sequences were annealed using reagents and protocols from the T4 polynucleotide kinase kit from New England Biolabs. The annealed gRNA sequences were then ligated into a BlpI/BstXI-digested plasmid containing a puromycin resistance gene from the Weissman laboratory (addgene #60955). The resulting plasmids were then cloned and amplified via heat shock transformation in competent DH5α bacteria. Bacteria suspensions were harvested and plasmids were isolated using a PureLink™ Quick Plasmid MiniPrep Kit (Invitrogen) and gRNA sequences were confirmed via next generation sequencing through Azenta Life Sciences. Purified plasmids (1 µg each) were transfected into HEK293T cells constitutively expressing dCas9-VPR using the Lipofectamine 3000 Transfection Reagent Kit (Thermo Fisher Scientific, Waltham, MA, USA) compared to an empty vector control. Following puromycin selection, cells were cultured for an additional three days and were subsequently lysed using TRIzol (Life Technologies, Carlsbad, CA, USA) for RNA purification.

**Table 1.**
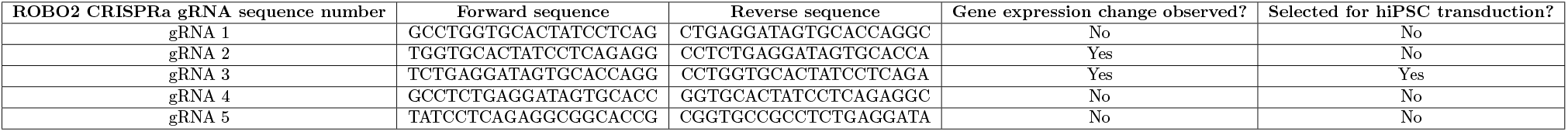
ROBO2 CRISPRa gRNA sequences and selection outcomes.

#### Lentivirus generation and hiPSC transduction

The gRNA plasmid that resulted in the greatest magnitude of ROBO2 gene activation was then selected for packaging into a lentivirus vector. Lentivirus was generated by transfecting HEK293T cells with the purified gRNA plasmid (3 µg), pMD2.G plasmid (0.6 µg, addgene #12259), and psPAX plasmid (1.4 µg, addgene #12260) using the Lipofectamine 3000 Transfection Reagent Kit. Cells were cultured in DMEM containing 10% FBS for the first 24 hours and were then switched to mTeSR plus medium for the following 72 hours. After this period, media was harvested and filtered and added to hiPSC cultured at *<*30% confluence in a 6-well plate and allowed to incubate for 24 hours. The following day, the media was switched to mTeSR plus containing puromycin (2 µg/mL) to select for hiPSCs transduced with the gRNA lentivirus.

#### RNA purification and RT-qPCR

Total RNA was purified from HEK293T cell culture using TRIzol purification followed by cDNA reverse transcription using the iScript kit (Biorad, Hercules, CA, USA). A 4 ng amount of cDNA was used in each 4 µL reaction using iTaq Universal SYBR® Green Supermix (Biorad, Hercules, CA, USA). All samples for all primers were plated in technical triplicates. Relative gene expression was determined using the 2^*−*!!*Ct*^ method, where genes of interest were normalized to the mean expression of two housekeeping genes (RPL27 and CYC1). qPCR primer sequences are listed below:

**Table 2.**
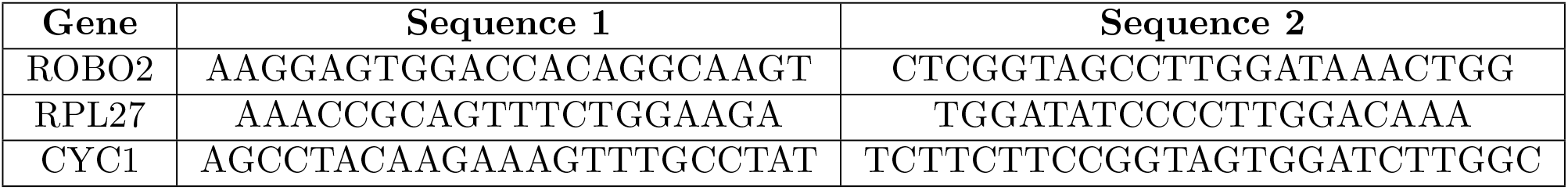
Primer sequences.

#### Western blot analysis

Neurons plated on 24-well plates and treated with doxycycline as described above were lysed in 100 µL of Tris lysis buffer (150 mM NaCl, 20 mM Tris pH 7.5, 1 mM EGTA, 1 mM EDTA, 1% Triton-X-100) and total protein abundance was measured by BCA assay kit (Thermo Scientific, Waltham, MA, USA). A 4.65 µg amount of total protein was loaded into a 4-15% gradient TGX gel (Bio-Rad) and transferred to a PVDF membrane (Bio-Rad). The membrane was then incubated with 4% paraformaldehyde for 30 minutes to fix proteins and was subsequently blocked with 5% nonfat dry milk in PBS for one hour at room temperature. The membrane was then incubated with primary antibody diluted in 5% bovine serum albumin in PBS overnight at 4ºC followed by horseradish peroxidase-conjugated secondary antibody diluted 1:7500 in 5% nonfat dry milk in PBS for one hour at room temperature. The membrane was then washed and treated with LICOR WesternSure PREMIUM Chemiluminescent substrate (Thermo Fisher Scientific) for 5 minutes followed by imaging on a Li-Cor Odyssey M imaging system (LI-COR Biosciences, Lincoln, NE, USA). *ROBO2* band intensity was normalized against GAPDH and all ROBO2/GAPDH ratios were normalized to the mean ROBO2/GAPDH ratios of the non-doxycycline treated lanes using ImageJ software. Antibody concentrations and dilutions are listed below:

**Table 3.** Primary antibodies and dilutions.

| **Protein Target** | **Manufacturer** | **Dilution** |
|---|---|---|
| ROBO2 | Cell Signaling Technology (Rabbit) | 1:1000 |
| GAPDH | Cell Signaling Technology (Rabbit) | 1:7500 |

#### Immunocytochemistry and Confocal Imaging

Neurons plated on 24-well plates and treated with doxycycline or vehicle control as described above were fixed with 4% PFA for 15 minutes then washed three times with PBS containing calcium and magnesium (PBS++). Cells were permeabilized with PBS++ containing 0.1% (v/v) Triton X-100 for 20 minutes, then washed twice with PBS++. Cells were then blocked with PBS++ containing 5% bovine serum albumin (BSA) at room temperature for 1 hour and incubated with primary antibodies against ROBO2 (rabbit) and MAP2 (chicken). All primary antibodies were diluted 1:500 in PBST (PBS++ containing 0.05% tween 20) containing 2.5% BSA. The following day, cells were washed three times with PBS++ followed by incubation with secondary antibodies (anti-rabbit Alexa Fluor 488 and anti-chicken Alexa Fluor 647) for 1 hour at room temperature. All secondary antibodies were diluted 1:500 in PBST containing 2.5% BSA. Cells were then washed three times with PBS++ and incubated with DAPI diluted in water (10 ng/mL) for 15 minutes. Cells were then mounted on glass slides with Prolong Gold Antifade mountant. Slides were imaged on a Leica TCS SP8 confocal laser microscope using a 63x/1.40 oil lens.

#### Total Tau and phospho-Tau analysis

Neurons plated on 24-well plates and treated with doxycycline as described above were lysed in 100 µL of Tris lysis buffer. Cell lysate was run on pTAUThr231/tTAU ELISA plates (Meso Scale Discovery) to quantify pTau and total Tau concentrations, respectively.

#### Statistical analysis

HEK293T validation data were analyzed using a two-tailed t-test in RStudio (R. Posit Software, PBC, Boston, MA). Tau and phospho-Tau data were analyzed using a two-tailed Welch’s t-test in GraphPad Prism (GraphPad Software Inc., La Jolla, CA, USA).

