## supplementary figures for "Disentangling Context-Specific Alzheimer’s Disease Progression Signatures through an Explainable AI Framework"

### Supplementary Appendix

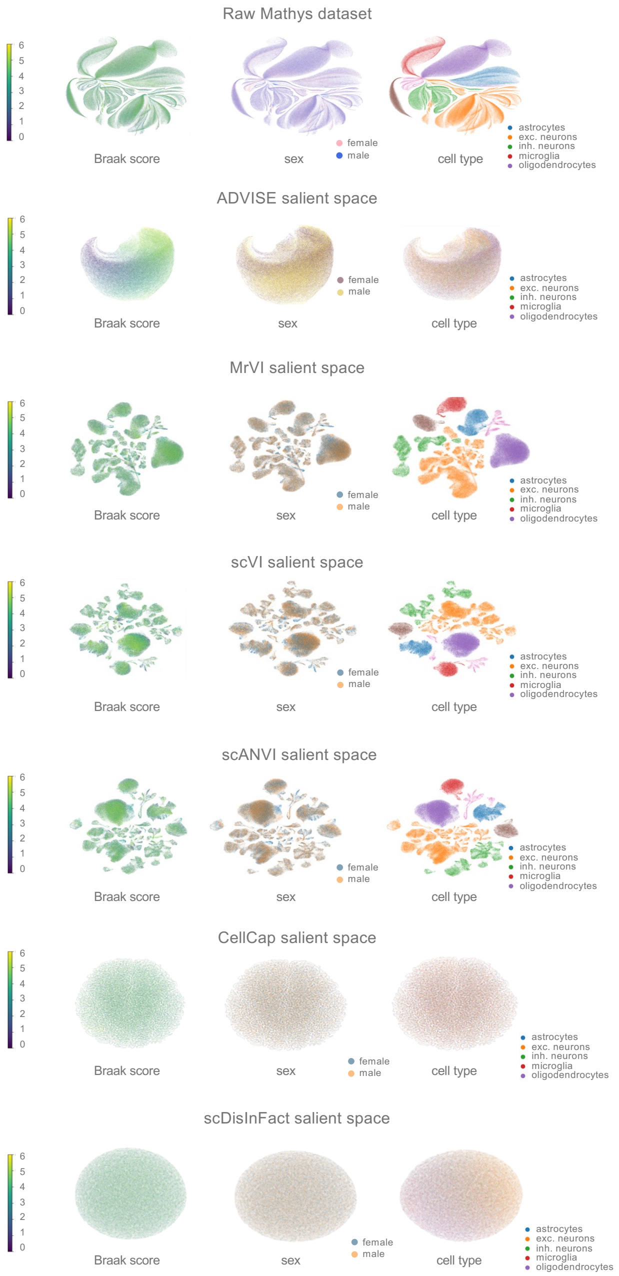


Supplementary Figure 1: **ADVISE achieves best Braak trend and background mixing.** The raw Mathys dataset is mainly separated by dominant cell type signatures. After ADVISE training, the salient space is able to capture a strong Braak trend with mixing in the covariate variables (cell type, sex).

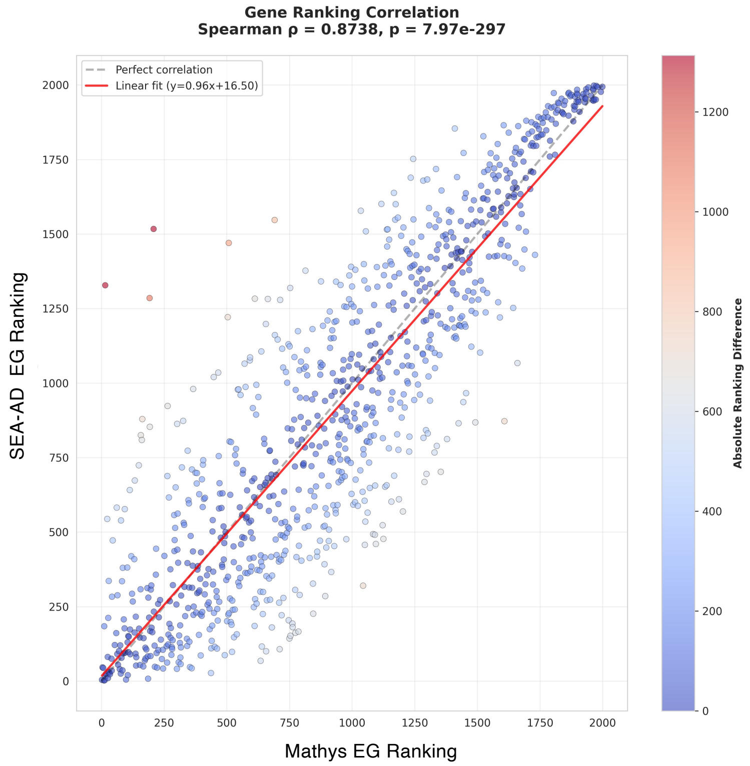


Supplementary Figure 2: **ADVISE achieves consistent gene rankings across AD datasets.** Among genes overlapping in the top 2,000 variable genes across both datasets, EG rankings from the Mathys-trained global model are plotted against those from the SEA-AD–trained model. Spearman correlation and significance indicate strong consistency between the two datasets.

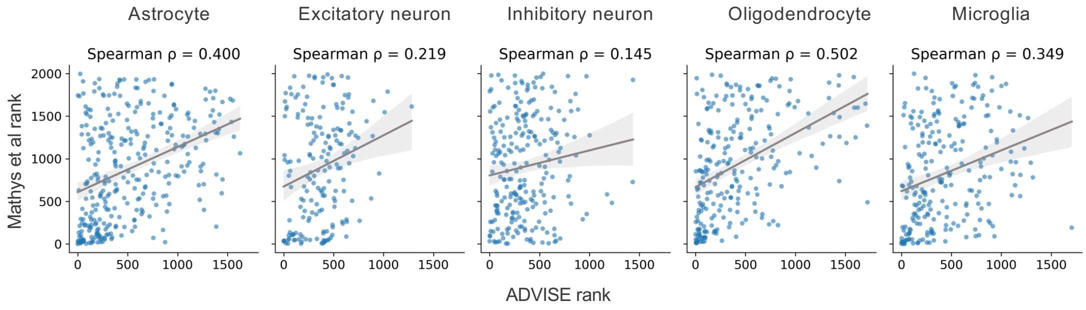


Supplementary Figure 3: **ADVISE uncovers new targets from original dataset.** For each cell type, rankings of genes overlapping in the top 2,000 lists were compared between ADVISE and the original Mathys et al. rankings. Overall, Spearman's correlation was low in neuronal populations and higher in glial cell types.

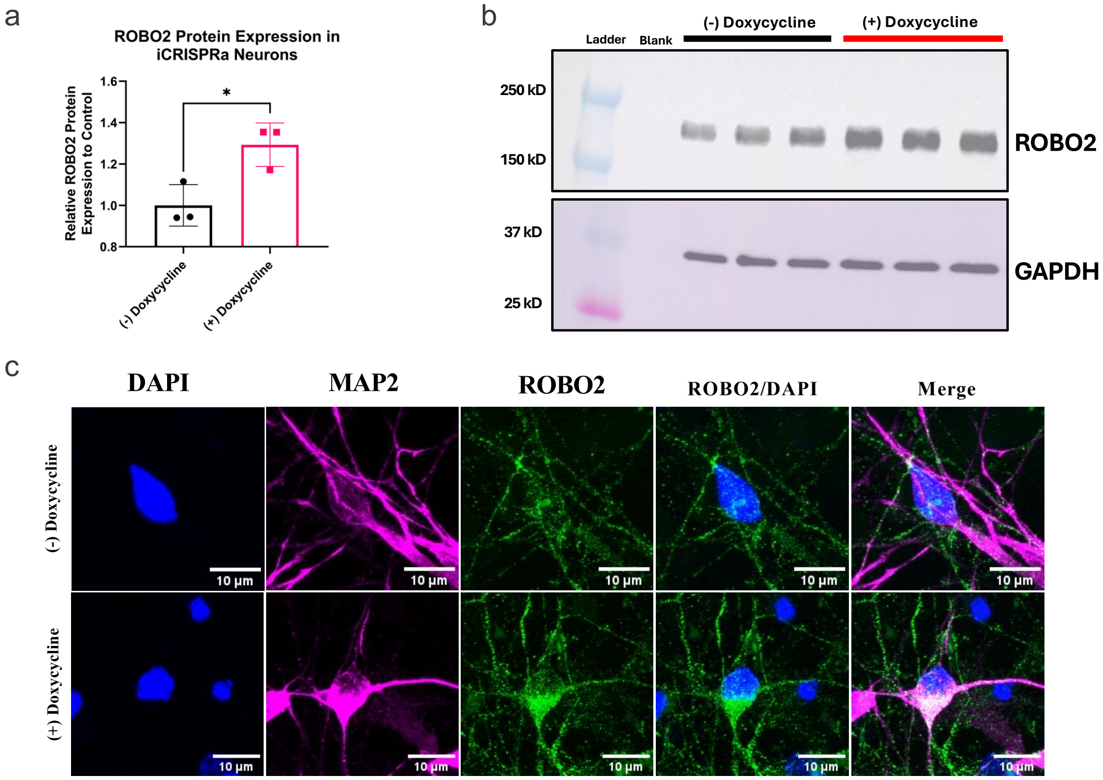


Supplementary Figure 4: ***ROBO2* protein expression measured by western blot and ICC images in hiPSC-derived neurons (a)** Quantification of western blot image. (**b**) *ROBO2* protein expression measured by western blot in hiPSC-derived neurons for neurons treated with doxycycline compared to a non-doxycycline treated control, with GAPDH being used as a loading control. *ROBO2* band intensity was normalized to GAPDH band intensity, all values were normalized to the mean ROBO2/GAPDH ratio of the non-doxycycline treated controls. Individual data points represent relative protein abundance values from independent lanes. (**c**) Representative ICC images of hiPSC-derived neurons with and without doxycycline treatment labeled with antibodies for *MAP2* (magenta) and *ROBO2* (green). Nuclei were counterstained with DAPI (blue).
